# Non-canonical stringent response signaling mediates antimicrobial fatty acid sensitivity in *Staphylococcus aureus*

**DOI:** 10.64898/2026.02.01.703180

**Authors:** Caitlin H. Kowalski, Sylvia B. Khalil, Dante A. James, T. Jarrod Smith, Killian D. Campbell, Rebecca M. Corrigan, Matthew F. Barber

## Abstract

*Staphylococcus aureus* is a leading cause of skin and soft tissue infections, yet colonization of healthy skin is limited by multiple defenses, including antimicrobial fatty acids (AFAs) produced by host cells and the resident microbiota. The mechanisms by which *S. aureus* overcomes these lipid-based defenses remain incompletely understood. Here, we show that mutations truncating the essential stringent response regulator Rel confer broad tolerance to both host-and microbially-derived AFAs in diverse *S. aureus* strains. Unlike classical stringent response activation, C-terminal Rel truncations do not induce typical (p)ppGpp-dependent transcriptional changes but instead enhance activity of the alternative sigma factor SigB and the staphylococcal accessory regulator SarA. This SigB-SarA regulatory cascade promotes transcriptional remodeling, including upregulation of pyrimidine biosynthesis genes, and coincides with alterations to cell envelope structure. Moreover, in the absence of SigB or SarA, tolerance can be restored through mutations in the serine/threonine phosphatase Stp1, highlighting additional pathways that modulate cell envelope-mediated resistance to AFAs. These findings identify a previously unrecognized consequence of Rel mutation in *S. aureus*; whereby small truncations may promote survival in AFA-rich host environments facilitating skin colonization and infection.

**Importance:** *Staphylococcus aureus* is a major cause of skin and soft tissue infections, and persistent skin colonization is a significant risk factor for infection. Antimicrobial fatty acids (AFAs), produced by microbes and host cells on human skin, normally limit *S. aureus* colonization. The mechanisms that allow this pathogen to overcome these lipid defenses are incompletely understood. We show that mutations truncating the essential stringent response regulator Rel enable *S. aureus* to tolerate both host-and microbially derived AFAs. These Rel variants, which have been identified in clinical isolates, alter cell envelope properties through the transcriptional regulator SarA rather than activating a classical stringent response. Our findings reveal a previously unrecognized adaptation that may facilitate *S. aureus* survival on the skin and promote infection.

## Introduction

In the healthcare setting, skin and soft tissue infections (SSTIs) are more common than urinary tract infections or pneumonia, with SSTI disease burden increasing in the United States (US) over the past decade (1, 2). *Staphylococcus aureus* is the pathogen most frequently associated with SSTIs and is estimated to cause about half of the SSTI hospitalizations in the US (3, 4). Skin colonization is a risk factor for infection, yet *S. aureus* colonization occurs only transiently or in low abundance at most healthy skin sites (5, 6). Innate skin defenses prevent *S. aureus* skin colonization including host antimicrobial peptides (7), antagonism by the resident microbiota (8), and the presence of antimicrobial fatty acids (AFAs) (9, 10). Understanding how *S. aureus* overcomes these defenses is essential for developing strategies to prevent *S. aureus*–associated SSTIs.

Diverse hosts, from plants to mammals, employ AFAs to directly inhibit bacterial pathogens (11, 12). On human skin, AFAs are produced by host and microbial lipolysis of stratum corneum or sebum lipid substrates (13–15). AFA potency is impacted by chain length, saturation, and the surrounding microenvironment(16, 17). While the mechanisms by which AFAs inhibit bacteria remain to be fully defined, they are capable of perturbing membranes (18, 19), causing lipid peroxidation (20), disrupting signaling pathways (21), and interfering with essential processes such as fatty acid synthesis and DNA replication (22, 23). *S. aureus* has evolved multiple strategies to combat AFAs including modulation of the cell surface (10), enzymatic detoxification (24, 25), and AFA efflux (26). Despite these mechanisms, *S. aureus* remains sensitive to AFA-rich human sebum and AFA application can successfully reduce *S. aureus* skin colonization in a murine model of atopic dermatitis (10). Additional mechanisms by which *S. aureus* adapts to tolerate AFAs remain to be fully defined.

We recently reported that the skin resident yeast *Malassezia sympodialis* generates the AFA 10-hydroxy palmitic acid (10-HP) that has potent bactericidal activity against *S. aureus* in skin-mimicking *in vitro* conditions (27). We isolated *S. aureus* mutant strains capable of tolerating 10-HP through repeated exposure to *M. sympodialis* cell-free supernatant (Ms-CFS) (**Figure S1A**). From four independent populations, in two strain backgrounds of Methicillin-Resistant *S. aureus* (MRSA), 10-HP tolerant strains acquired independent mutations in the stringent response regulator Rel. Two Rel mutations investigated further, L127V and Q672*, were sufficient for 10-HP tolerance suggesting a role for the stringent response in mediating AFA sensitivity in *S. aureus.* The stringent response (SR) is a highly conserved stress response that is classically induced by amino acid starvation but also responds to cell-wall damage and alkaline conditions, among other stressors (28). SR is orchestrated by synthesis and degradation of the intracellular phosphorylated nucleotide messengers guanosine pentophosphate and guanosine tetraphosphate (pppGpp and ppGpp, respectively), collectively called (p)ppGpp or alarmones. (p)ppGpp are synthesized by one or multiple GTP pyrophosphokinases from GTP or GDP (29). In Pseudomonadota (previously Proteobacteria), (p)ppGpp interacts directly with RNA polymerase to regulate transcription resulting in reduced growth rate and altered metabolism that facilitates survival in unfavorable conditions (29, 30). In Bacillota (previously Firmicutes), (p)ppGpp does not directly bind RNA polymerase but instead controls GTP levels that impact nucleotide sensitive promoters and GTP-sensing transcriptional regulators to control survival and growth (31, 32). *S. aureus* encodes three proteins that synthesize (p)ppGpp: Rel, RelQ, and RelP. Where RelQ and RelP are synthetase-only GTP pyrophosphokinases, Rel is a dual-function protein that synthesizes and degrades (p)ppGpp (29). Rel hydrolase activity renders the protein conditionally essential in *S. aureus*, as complete loss of hydrolase activity in the presence of RelP and RelQ synthetase activity stalls bacterial growth (33). Despite its essential functions, recent studies have identified natural Rel variants in *S. aureus* clinical isolates that alter bacterial tolerance to clinical antibiotics and nutrient stress (34–36). This suggests that mutations in Rel tune the SR to permit *S. aureus* to overcome unfavorable host conditions, however the role of the SR in broad AFA tolerance has not been investigated.

Here, we leverage a collection of naturally occurring and laboratory derived Rel allele variants to investigate the impact of Rel mutations on AFA sensitivity in *S. aureus.* These variants include both missense and nonsense mutations across the coding sequence within both the N-terminal catalytic domain and C-terminal regulatory domain (**Figure 1A**). We further investigate the mechanisms through which truncation of Rel impacts AFA sensitivity, utilizing 10-HP and the polyunsaturated fatty acid linoleic acid as representatives of microbial or host derived AFAs, respectively. We report that Rel truncation, by as little as 57 amino acids, reduces *S. aureus* AFA sensitivity through increased activity of the staphylococcal accessory regulator SarA. Notably, C-terminal truncation of Rel does not appear to activate the canonical SR transcriptional response, but results in changes to the cell envelope that coincide with AFA tolerance. Together these data support a model by which C-terminal truncation of the essential protein Rel can occur during host colonization or infection resulting in strains capable of tolerating microbial and host AFAs.

**Figure 1.**
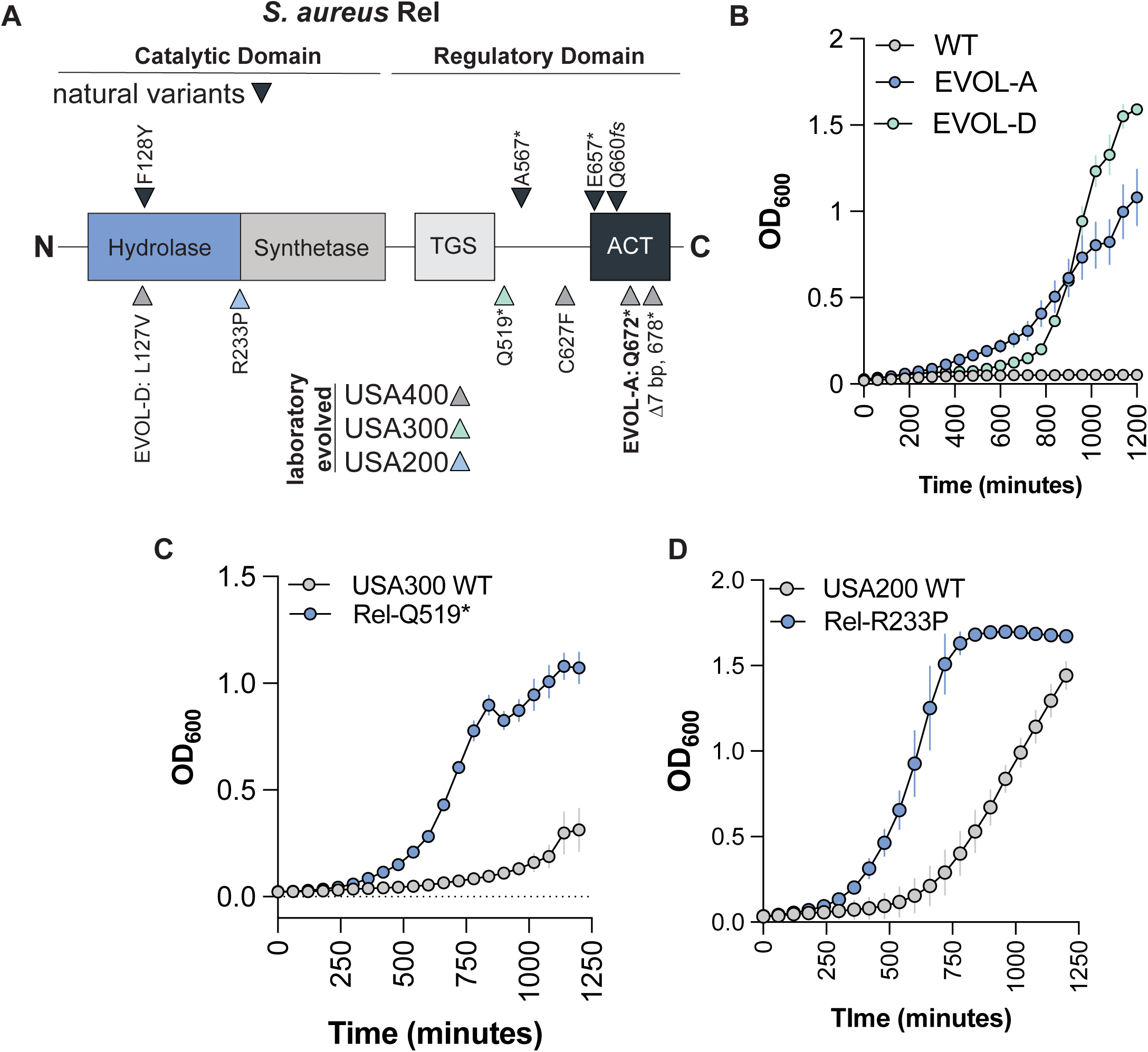
Mutations in Rel across diverse strains coincide with decreased sensitivity to the antimicrobial fatty acid linoleic acid. A. Schematic of Rel protein domains and mutations that occurred during laboratory evolution or identified in natural variants. B-D. 20-hour growth curve in TSB with 112 µg/mL linoleic acid showing mean OD_600_ from three separate experiments.

## Results

### Mutations in the stringent response regulator Rel reduce linoleic acid sensitivity in multiple strain lineages

We previously isolated *S. aureus* USA300 and USA400 strains with resistance to the AFA 10-HP produced by *M. sympodialis* following repeated exposure to Ms-CFS (**Figure S1A**)(27). The USA400 strain, NRS193, is a clinical isolate from a bacteremia infection and is closely related to the USA400 reference strain MW2 (**Table S1**). In this study, NRS193 is referred to as the wild type (WT) strain. In this strain background, we isolated independent Rel mutations following repeated exposure to Ms-CFS (Rel-C627F, Rel-Q672*, Rel-678*) or long-term co-culture with *M. sympodialis* (Rel-L127V) (**Figure 1A**). We named the N-terminal Rel-L127V strain EVOL-D and the Rel-Q672* strain EVOL-A, and found both strains to be tolerant to Ms-CFS and purified 10-HP(27). To determine if these strains are broadly tolerant to AFAs, we used the model host-derived polyunsaturated AFA linoleic acid. Where WT (NRS193) growth was completely inhibited by 112 µg/mL linoleic acid, the Rel mutant strains EVOL-A and EVOL-D were able to reach terminal optical densities (OD_600_) greater than 1 (**Figure 1B**). Similarly, in a USA300 strain background (**Table S1**), a Ms-CFS adapted strain with a Rel mutation (Rel-Q519*) was found to be tolerant to Ms-CFS/10HP (27) and to have increased growth in the present of linoleic acid (**Figure 1C**). Together these data suggest that Rel mutations in MRSA lineages can confer reduced sensitivity to linoleic acid in addition to 10-HP.

The SR has been linked to homogenous resistance to β-lactams in MRSA strains through modulation of *mecA* gene expression (37). Given that the cell envelope is the initial interaction site of AFAs with the cell, we sought to determine if methicillin sensitive *S. aureus* (MSSA) would also adapt to AFAs via mutation of Rel. Through the same repeated exposure to Ms-CFS with the MSSA USA200 reference strain MN8 (**Table S1**), we observed rapid adaptation after 12 passages (**Figure S1B**). We isolated a strain from the terminal passage with increased tolerance to Ms-CFS (**Figure S1C**) and 10-HP (**Figure S1D**). Whole genome sequencing revealed this strain had a Rel mutation in the N-terminal hydrolase domain (Rel-R233P) (**Figure 1A**). Similarly to the USA300 and USA400 strains, the mutation of Rel in USA200 also conferred increased growth in the presence of the AFA linoleic acid (**Figure 1D**). From these data, we conclude that mutations in Rel can tune AFA sensitivity across diverse *S. aureus* strains independent of methicillin resistance.

### Rel mutations are sufficient for reduced sensitivity to structurally diverse AFAs

*S. aureus* is sensitive to structurally distinct AFAs including saturated fatty acids, monounsaturated fatty acids, and polyunsaturated fatty acids (38) (**Figure 2A**). We selected lauric acid, palmitoleic acid, and linoleic acid to represent each class, respectively, and sought to determine if Rel mutations were sufficient to modulate sensitivity to each AFA. As shown for linoleic acid in **Figure 1**, different *S. aureus* strains display inherent differences in AFA sensitivity. To overcome this, we leveraged our previously developed collection of NRS193 strains where the WT Rel allele was replaced with Rel variants from the lab (Rel-Q672* and Rel-L127V) or Rel variants from naturally occurring strains (Rel-F128Y, Rel-E657*, Rel-A567*, Rel-Q660*fs*) (**Figure 1A, Table S1**)(27). The natural variants are from clinical isolates or identified by querying publicly available *S. aureus* genomes (35).

**Figure 2.**
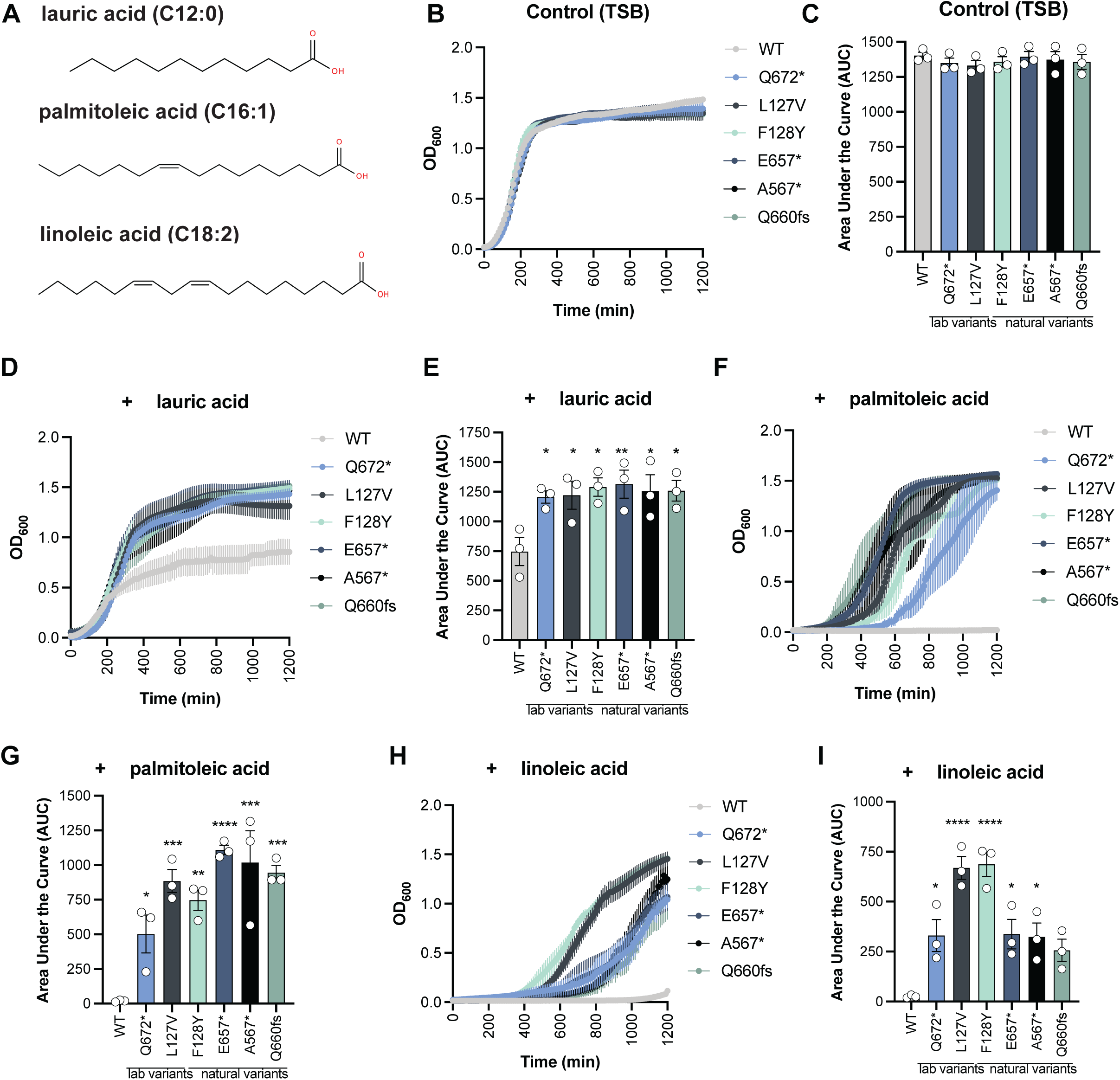
Mutant Rel alleles are sufficient to reduce sensitivity to diverse antimicrobial fatty acids. A. Structures of the three antimicrobial fatty acids lauric acid, palmitoleic acid, and linoleic acid B. 20-hour growth curve in TSB showing mean OD_600_ from three separate experiments. C. AUC calculated from growth curves in A. Each point represents a separate experiment. D. 20-hour growth curve in TSB with 75 µg/mL lauric acid showing mean OD_600_ from three separate experiments. E. AUC calculated from growth curves in C. Each point represents a separate experiment. F. 20-hour growth curve in TSB with 37.5 µg/mL palmitoleic acid showing mean OD_600_ from three separate experiments. G. AUC calculated from growth curves in E. Each point represents a separate experiment. H. 20-hour growth curve in TSB with 112 µg/mL linoleic acid showing mean OD_600_ from three separate experiments. I. AUC calculated from growth curves in G. Each point represents a separate experiment. *:p<0.05, **:p<0.01, ***:p<0.001, ****p<0.0001, ns: p>0.05

We cultured the WT (NRS193) and the Rel mutants in the standard laboratory media tryptic soy broth (TSB) as the control condition. We monitored cultures over 20 hours and quantified growth as the area under the curve (AUC). The overall growth in TSB was not significantly different for any of the mutants compared to WT (**Figure 2B, C**). In the presence of the saturated AFA lauric acid (75 µg/mL), all Rel alleles, whether laboratory variants or natural variants, were sufficient to significantly increase growth compared to WT (**Figure 2D, E**). Similarly, all Rel variants showed significantly increased growth in the presence of the monounsaturated AFA palmitoleic acid (37.5 µg/mL) (**Figure 2F, G**). Notably, while the Rel variants, apart from Rel-Q660*fs*, grew significantly better in the presence of the polyunsaturated AFA linoleic acid (112 µg/mL), strains with mutations in the hydrolase domain (Rel-L127V and Rel-F128Y) were phenotypically distinct from the C-terminal variants (**Figure 2H, I**). The hydrolase Rel mutants have a shorter lag phase and reach a higher terminal OD_600_ compared to the other Rel variants perhaps due to increased (p)ppGpp production. Together these data suggest that the Rel-mediated AFA tolerance is broad enough to apply to structurally distinct AFAs and that the specific mutations can differentially tune AFA sensitivity. We next sought to identify the mechanisms downstream of Rel that mediate the response to AFAs.

### The C-terminal Rel mutant, EVOL-A, does not trigger classical SR activation

The best characterized Rel variant is the hydrolase mutant Rel-F128Y. This variant was identified from a case of persistent bacteremia and is sufficient for tolerance to diverse clinical antibiotics. The mutation in the hydrolase domain reduces hydrolase function and results in elevated (p)ppGpp which reduces *S. aureus* growth, ultimately leading to antibiotic tolerance (35, 36). Consistent with this model, we have also previously observed increased (p)ppGpp in *S. aureus* NRS193 carrying the hydrolase mutant Rel-L127V compared to WT after SR induction with mupirocin(27). Notably, the same (p)ppGpp increase was not evident with the C-terminal mutant Rel-Q672* compared to WT (27). This suggest that the downstream mechanisms leading to AFA tolerance may also depend on the specific Rel mutation, since GDP/GTP consumption to synthesize (p)ppGpp is what largely drives the SR-dependent transcriptional changes in *S. aureus*.

C-terminal Rel mutations have also been identified in *S. aureus* clinical isolates and based on our data these mutations, in a range of 57 to 162 amino acids, can impact AFA sensitivity (**Figure 2**) (27, 34). As the consequences of these C-terminal mutations are less clear compared to the Rel hydrolase mutations, we focused on the EVOL-A (Rel-Q672*) mutant to investigate downstream AFA tolerance mechanisms. We first performed RNA-sequencing to understand the basal differences in the transcriptome prior to AFA exposure when strains were grown in TSB. Analysis of differentially expressed genes between EVOL-A and WT (NRS193) did not reveal robust SR activation in EVOL-A. In *S. aureus*, SR transcriptional changes are largely orchestrated by the transcriptional repressor CodY that responds to intracellular changes in GTP and branched chain amino acids to control expression of central metabolism genes (39). There was no clear evidence of CodY de-repression in EVOL-A, with no significant changes in the representative CodY regulated genes *ilvD, leuA,* or *metH* (**Table 1**). This supports our previous observation that CodY does not play a role in 10-HP tolerance in NRS193(27). CodY-independent changes also occur during SR activation (40). While not characterized in a USA400 background specifically, these include induction of the phenol-soluble modulins (*psmA1-4* and *psmB1-2*) and repression of the *gua* operon (*guaA, guaB, hpt, pbuX*) (39). In EVOL-A compared to WT, we see modest but significant changes in *psmA2-4* and the *gua* operon consistent with previous results during SR induction, but at a magnitude for which the biological significance is unclear (**Table 1**). Based on these results, we conclude that the C-terminal truncation of Rel does not lead to robust classic SR activation in *S. aureus*.

**Table 1.**
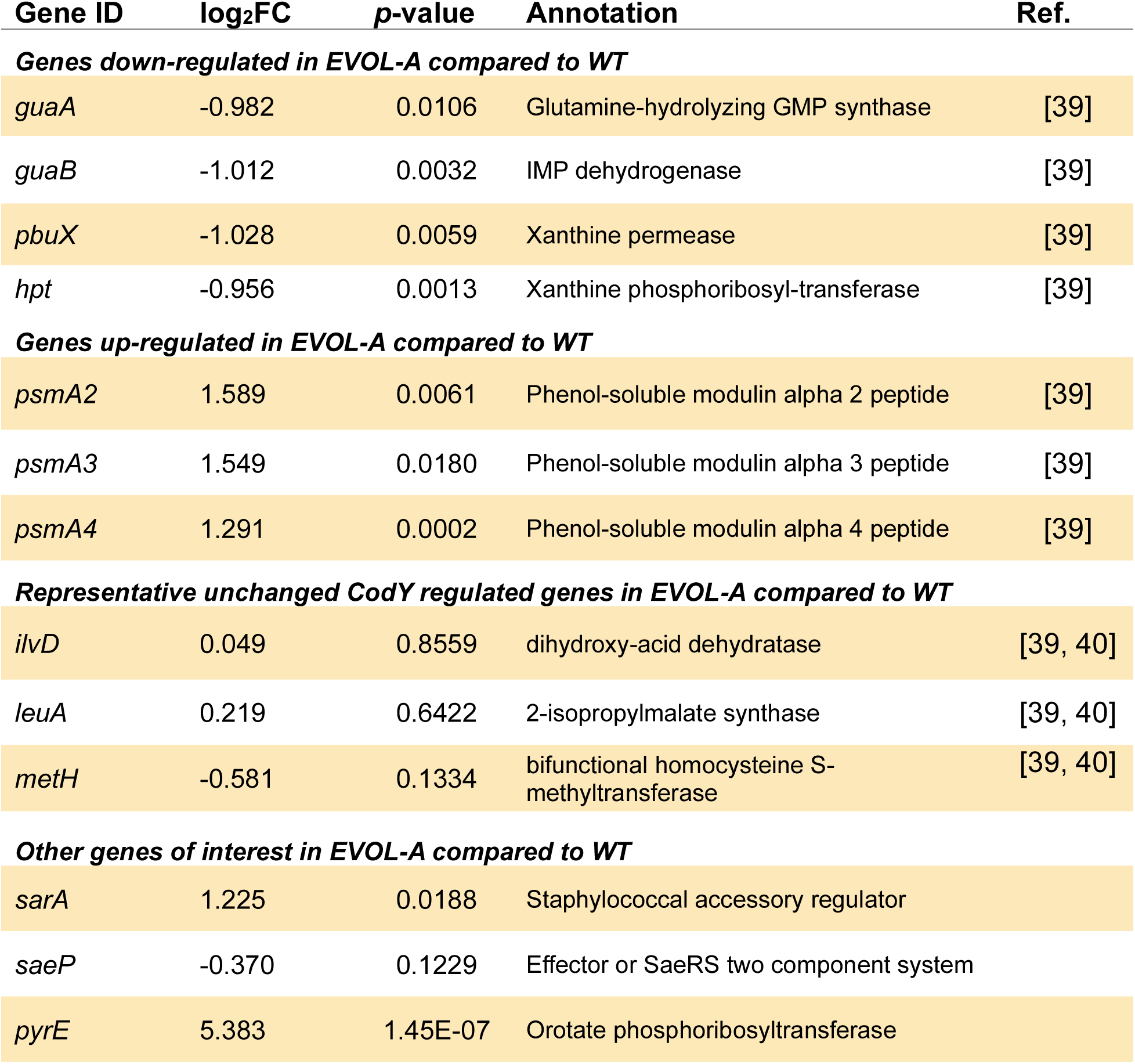
Selected differentially expressed genes in EVOL-A compared to WT.

| Gene ID | log<sub>2</sub>FC | p-value | Annotation | Ref. |
| --- | --- | --- | --- | --- |
| ***Genes down-regulated in EVOL-A compared to WT*** |  |  |  |  |
| *guaA* | -0.982 | 0.0106 | Glutamine-hydrolyzing GMP synthase | [39] |
| *guaB* | -1.012 | 0.0032 | IMP dehydrogenase | [39] |
| *pbuX* | -1.028 | 0.0059 | Xanthine permease | [39] |
| *hpt* | -0.956 | 0.0013 | Xanthine phosphoribosyl-transferase | [39] |
| ***Genes up-regulated in EVOL-A compared to WT*** |  |  |  |  |
| *psmA2* | 1.589 | 0.0061 | Phenol-soluble modulins alpha 2 peptide | [39] |
| *psmA3* | 1.549 | 0.0180 | Phenol-soluble modulins alpha 3 peptide | [39] |
| *psmA4* | 1.291 | 0.0002 | Phenol-soluble modulins alpha 4 peptide | [39] |
| ***Representative unchanged CodY regulated genes in EVOL-A compared to WT*** |  |  |  |  |
| *ilvD* | 0.049 | 0.8559 | dihydroxy-acid dehydratase | [39, 40] |
| *leuA* | 0.219 | 0.6422 | 2-isopropylmalate synthase | [39, 40] |
| *metH* | -0.581 | 0.1334 | bifunctional homocysteine S-methyltransferase | [39, 40] |
| ***Other genes of interest in EVOL-A compared to WT*** |  |  |  |  |
| *sarA* | 1.225 | 0.0188 | Staphylococcal accessory regulator |  |
| *saeP* | -0.370 | 0.1229 | Effector or SaeRS two component system |  |
| *pyrE* | 5.383 | 1.45E-07 | Orotate phosphoribosyltransferase |  |

### The staphylococcal accessory regulator SarA controls the transcriptional profiles of EVOL-A

We next sought to determine how Rel C-terminal truncation could result in broad AFA tolerance if not through classical activation of the SR. While the transcriptome data did not reveal the CodY-dependent changes characteristic of the SR, there were 148 significant differentially expressed genes between EVOL-A and WT (NRS193) (**Table S2**). Of these, 60 genes were downregulated in EVOL-A and 88 genes were upregulated in EVOL-A compared to WT. Most notably, genes involved in *de novo* pyrimidine biosynthesis were upregulated in EVOL-A (**Figure 3A**, **Tables S2**). In addition, *S. aureus* Protein A (*spa*) and a predicted operon involved in amino acid metabolism (*ald>tdcB>potE>norB)*, corresponding to MW1325-MW1328 in the USA400 reference strain MW2, were significantly downregulated in EVOL-A compared to WT in baseline conditions (**Figure 3A**).

**Figure 3.**
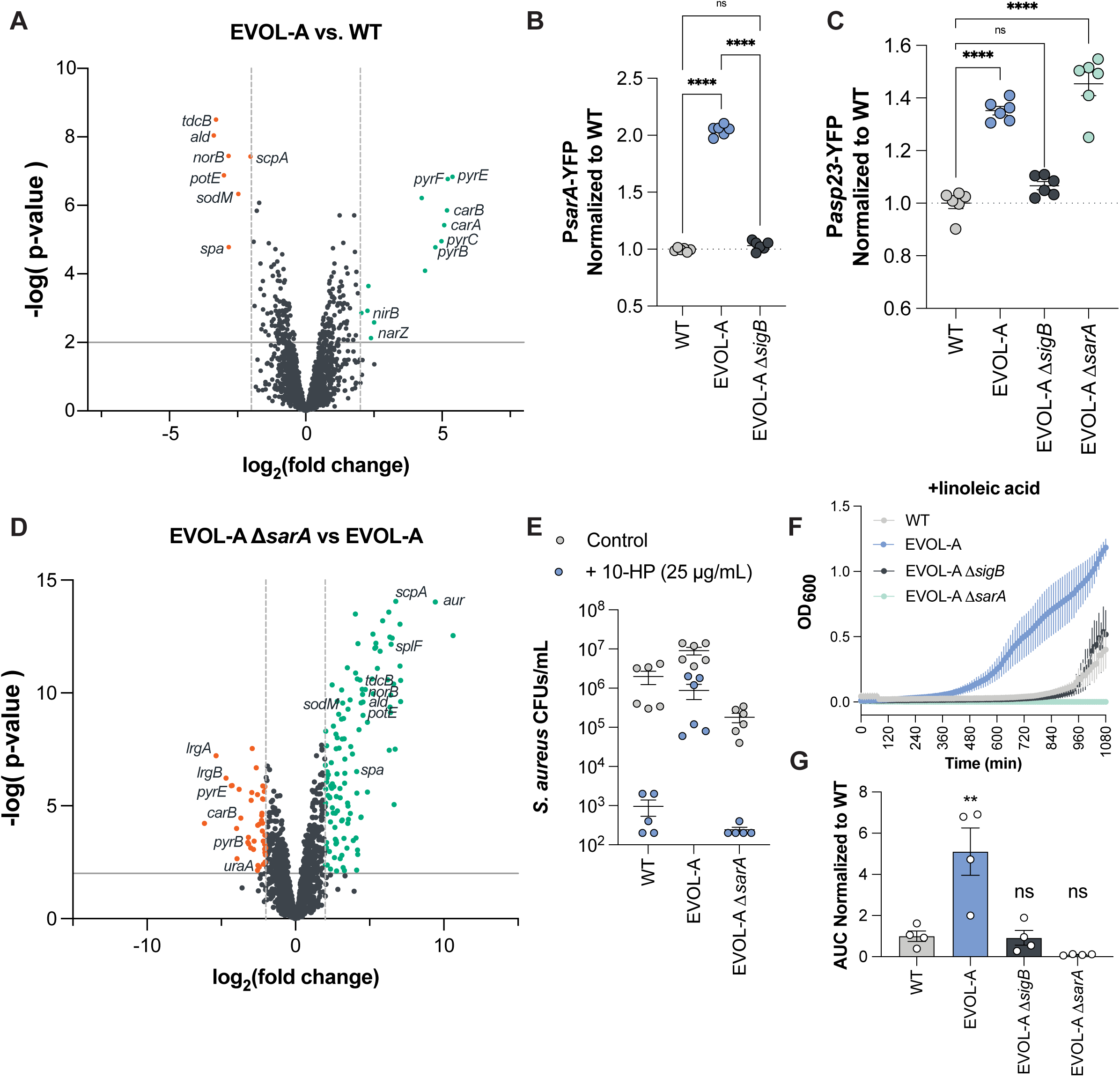
The accessory regulator SarA shapes the transcriptome of EVOL-A and is necessary for antimicrobial fatty acid tolerance. A. Volcano plot displaying differentially expressed genes in the EVOL-A strain compared to WT in baseline conditions. Green indicates transcripts increased in EVOL-A and orange indicate transcripts decreased in EVOL-A compared to WT. B. Transcriptional reporter expression for *sarA* promoter across strains and normalized to WT. C. Transcriptional reporter expression for SigB activity using the promoter of *asp23* across strains and normalized to WT. D. Volcano plot displaying differentially expressed genes in the EVOL-A Δ*sarA* strain compared to EVOL-A. E. Recovered colony forming units (CFUs) after treatment with 10-HP or the control condition for 4 hours. Data pooled from two experiments. F. 18-hour growth curve in TSB with 112 µg/mL linoleic acid showing mean OD_600_ from four separate experiments. G. AUC calculated from growth curves in F. Each point represents a separate experiment. *:p<0.05, **:p<0.01, ***:p<0.001, ****p<0.0001, ns: p>0.05

We had previously shown using a transcriptional reporter that the activity of the alternative sigma factor SigB is increased in EVOL-A compared to WT, and that SigB is essential for 10-HP tolerance in EVOL-A (27). However, RNA-sequencing at this time point did not reveal significant changes in the SigB reporter gene *asp23* (**Table S3**). We sought to understand how the transcriptional changes in EVOL-A at baseline were controlled to determine how the C-terminal Rel truncations impact EVOL-A physiology and AFA tolerance. The downregulation of *spa* and the cysteine protease *scpA* in EVOL-A compared to WT suggested that the staphylococcal accessory regulator SarA might contribute to the expression pattern of EVOL-A (**Figure 3A**, **Table S2**). SarA is a RNA and DNA-binding protein that regulates the transcription of genes and transcript stability in *S. aureus* and is a well-characterized repressor of the virulence factor *spa* and proteases including *scpA* (41, 42). *sarA* expression was also modestly but significantly increased in EVOL-A compared to WT in the RNA-sequencing data (**Table S2**). To confirm this, we generated a transcriptional reporter for *sarA* expression by encoding yellow fluorescent protein (YFP) downstream of the full *sarA* promoter (P1, P2, P3) and introduced it at a neutral site on the chromosome of WT (NRS193) and EVOL-A (43). We observed a similar, approximately 2-fold increase in *sarA* expression in EVOL-A compared to WT as seen in the RNA-sequencing data (**Figure 3B**).

To determine if SarA could be acting downstream of the elevated SigB activity that we previously reported, we assessed SarA promoter activity in EVOL-A in the absence of SigB (27). There is conflicting literature, across different *S. aureus* strains, as to whether SigB regulates SarA (44). Our data suggest that the elevated *sarA* expression in EVOL-A is SigB-dependent as in the absence of *sigB, sarA* promoter activity is significantly reduced (**Figure 3B**). To further investigate the model that the Rel C-terminal truncation in EVOL-A activates SigB which in turn activates SarA, we generated a *sarA* deletion mutant in EVOL-A to determine if SigB activity was altered. We utilized the well-characterized *asp23* promoter as a reporter for SigB activity(45). As previously observed, SigB activity is modestly but significantly increased in EVOL-A, and loss of *sarA* does not impact SigB activity (**Figure 3C**)(27). These data support the model whereby SigB acts upstream to regulate *sarA*.

To confirm our hypothesis that elevated SarA activity drives the transcriptional profile of the C-terminal Rel mutant strain EVOL-A, we performed RNA-sequencing to compare EVOL-A and EVOL-A Δ*sarA*. Consistent with the role of SarA as a master regulator of *S. aureus* transcription, there were 523 significant differentially expressed genes in EVOL-A compared to EVOL-A Δ*sarA* (**Table S4**). This included 214 genes with significant reduced expression in EVOL-A Δ*sarA* and 309 with significant increased expression. Consistent with *sarA* deletion, *spA* and protease (*scpA*, *aur*) transcripts were significantly increased in abundance in the absence of *sarA* (**Figure 3D**). More surprising was the apparent SarA-dependent expression of pyrimidine metabolism in EVOL-A, as well as SarA-dependent repression of the putative amino acid metabolism operon *ald>tdcB>potE>norB.* The latter group of genes includes a putative alanine dehydrogenase (ald), a putative threonine ammonia-lyase (tdcB), an MFS transporter (norB), and a putative amino acid permease (potE) (**Table S3**). While the regulation of these pathways by SarA is not well described, there are reports in the literature where loss of *sarA* results in suppression of pyrimidine biosynthesis (46) and at least one example of SarA-dependent regulation of *tdcB* (also called *ilvA1*) (47). Together, these data indicate that in the context of a C-terminal Rel truncation (EVOL-A), the transcriptional state of the cell is largely controlled by increased SarA activity. We next sought to determine if SarA contributes to the broad AFA tolerance in the context of Rel truncation.

### SigB and SarA are essential for tolerance to AFA in the Rel mutant strain EVOL-A

We first assessed the contribution of SarA to 10-HP tolerance in EVOL-A. We’ve previously shown that *sigB* is necessary for 10-HP tolerance in EVOL-A and in the context of another C-terminal Rel truncation mutant: Rel-E657* (27, 48). Consistent with SarA acting downstream of SigB, Δ*sarA* abolishes 10-HP tolerance in EVOL-A (**Figure 3E**). We next assessed broader AFA tolerance using the model polyunsaturated AFA linoleic acid. As previously shown, EVOL-A grows significantly better than WT in the presence of linoleic acid. This enhanced growth is dependent on both *sigB* and *sarA* (**Figure 3F**). In the context of the other C-terminal Rel truncation mutant Rel-E657*, *sigB* and *sarA* also contribute to reduced 10-HP and linoleic acid sensitivity (**Figure S2**).

We hypothesized that the SarA dependent transcripts differentially expressed in EVOL-A compared to WT would contribute to AFA tolerance. As an initial screen, we utilized the Nebraska Transposon Mutant Library (NTML) generated in the MRSA JE2 strain background(48). We selected available transposon insertion mutants for genes with reduced expression in the EVOL-A compared to WT (*tdcB*::Tn, *ald*::Tn, *potE*::Tn, and *spA*::Tn). We hypothesized that downregulation of these genes might contribute to AFA tolerance. However, none of the transposon mutants displayed increased tolerance to 10-HP (**Figure S3A**). To determine how SarA and SigB mediate AFA tolerance in EVOL-A, we decided to apply a forward genetics approach.

### Evolution of AFA tolerance in the absence of SigB or SarA implicates the serine/threonine phosphatase Stp1

We hypothesized that we could select to restore AFA tolerance in EVOL-A lacking either *sigB* or *sarA*, and that this would allow us to identify the downstream mechanisms of tolerance. To do this, we applied a similar experimental evolution strategy used to generate EVOL-A by serially exposing EVOL-A Δ*sigB* or EVOL-A Δ*sarA* to Ms-CFS (**Figure S4A, D**). For both strains, we observed rapid enrichment for Ms-CFS tolerance after only three to five passages across independent populations (**Figure S4B, E**). Isolation of single colonies from the final evolved populations yielded strains with stable tolerance to Ms-CFS (**Figure S4C, F**). Many isolates were as tolerant as EVOL-A, particularly for those generated in the EVOL-A Δ*sarA* background (**Figure S4F**). We selected seven isolates from the EVOL-A Δ*sigB* experiment and five isolates from the EVOL-A Δ*sarA* experiment for whole genome sequencing and variant analysis (**Table 2**). For EVOL-A Δ*sigB* evolved isolates, the Replicate 1 (R1) population had a missense mutation in the essential sensor histidine kinase *walK.* The R3 population had isolates with three missense mutations in the anaerobic redox response regulator *airR*, the acetyl-CoA carboxylase subunit *accD*, and the serine/threonine phosphatase *stp1*. R3 also had an isolate with only a nonsense mutation in the mannose-6-phosphate isomerase *manA*. For EVOL-A Δ*sarA* evolved isolates, the R1 population has a missense mutation in a purine biosynthetic gene *purQ*. The R2 population has isolates with a frameshift mutation in the serine/threonine phosphatase *stp1* both alone and in combination with a missense mutation in the chaperonin *groL*. The R3 isolate had the same frameshift mutation in *stp1* as R2. The enrichment for *stp1* mutations in independent populations in both strains encouraged us to follow up on its potential role in AFA tolerance.

**Table 2.**
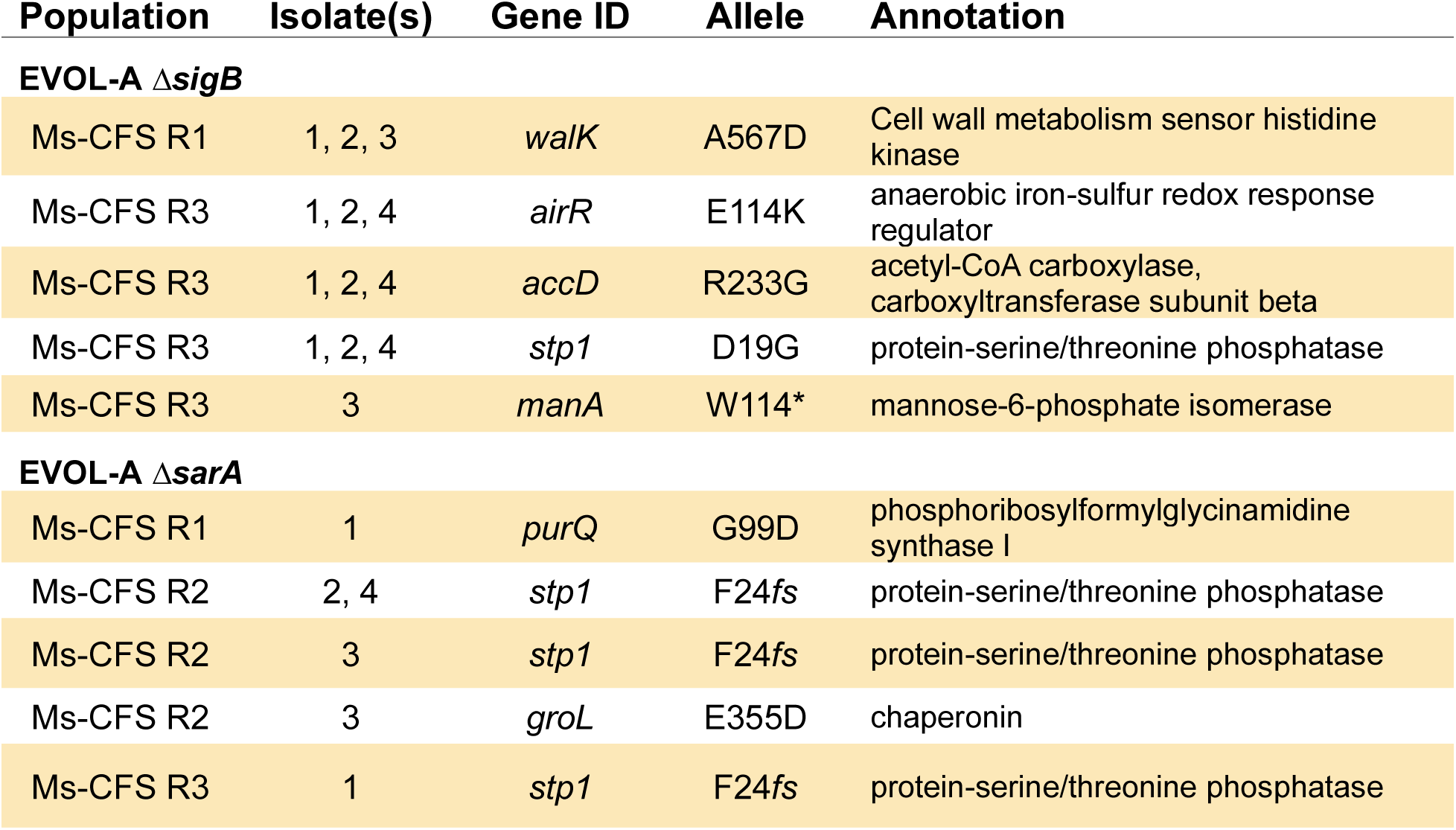
Variant analysis from evolved Ms-CFS tolerant isolates in EVOL-A Δ*sigB* and EVOL-A Δ*sarA* ancestral strains.

### Loss of function mutations in stp1 reduce AFA sensitivity in WT and Rel mutant strains

The *stp1* mutation that arose in EVOL-A *ΔsarA* (Stp1-F24*fs*) causes a frameshift at amino acid 24 of 247 and is expected to cause loss of function. The *stp1* mutation that arose in EVOL-A Δ*sigB* is a missense mutation that changes an aspartate to a glycine (D19G) and is not as clearly a loss of function mutation. In EVOL-A, the *stp1* mutation also occurs with two other mutations and is thus not clearly the causative mutation of the AFA/Ms-CFS tolerance. To address these questions, we introduced the evolved *stp1* mutant allele (*stp1*^D19G^) or deleted the entire *stp1* coding sequence (Δ*stp1*) in EVOL-A Δ*sigB* and assessed Ms-CFS, 10-HP, and linoleic acid sensitivity. Both the mutant allele and loss of *stp1* in EVOL-A Δ*sigB* restored Ms-CFS (**Figure S4G**) and 10-HP tolerance comparable to EVOL-A (**Figure 4A**) and significantly increased growth in the presence of linoleic acid (**Figure 4B, C**). Stp1 is the cognate phosphatase to the serine/threonine kinase Stk1 (also called PknB) (49). Since loss of Stp1 facilitates 10-HP tolerance in EVOL-A Δ*sigB*, we hypothesized that tolerance would be dependent on the Stk1 kinase. We generated a *stk1* deletion in the EVOL-A Δ*sigB* +*stp1*^D19G^ strain and observed that without *stk1*, 10-HP tolerance is reduced ∼1,000-fold (**Figure 4A**). Together these data suggest that in the absence of *sigB* and *sarA*, increased Stk1 kinase activity, mediated by reduced Stp1 phosphatase activity, is sufficient to restore AFA tolerance.

**Figure 4.**
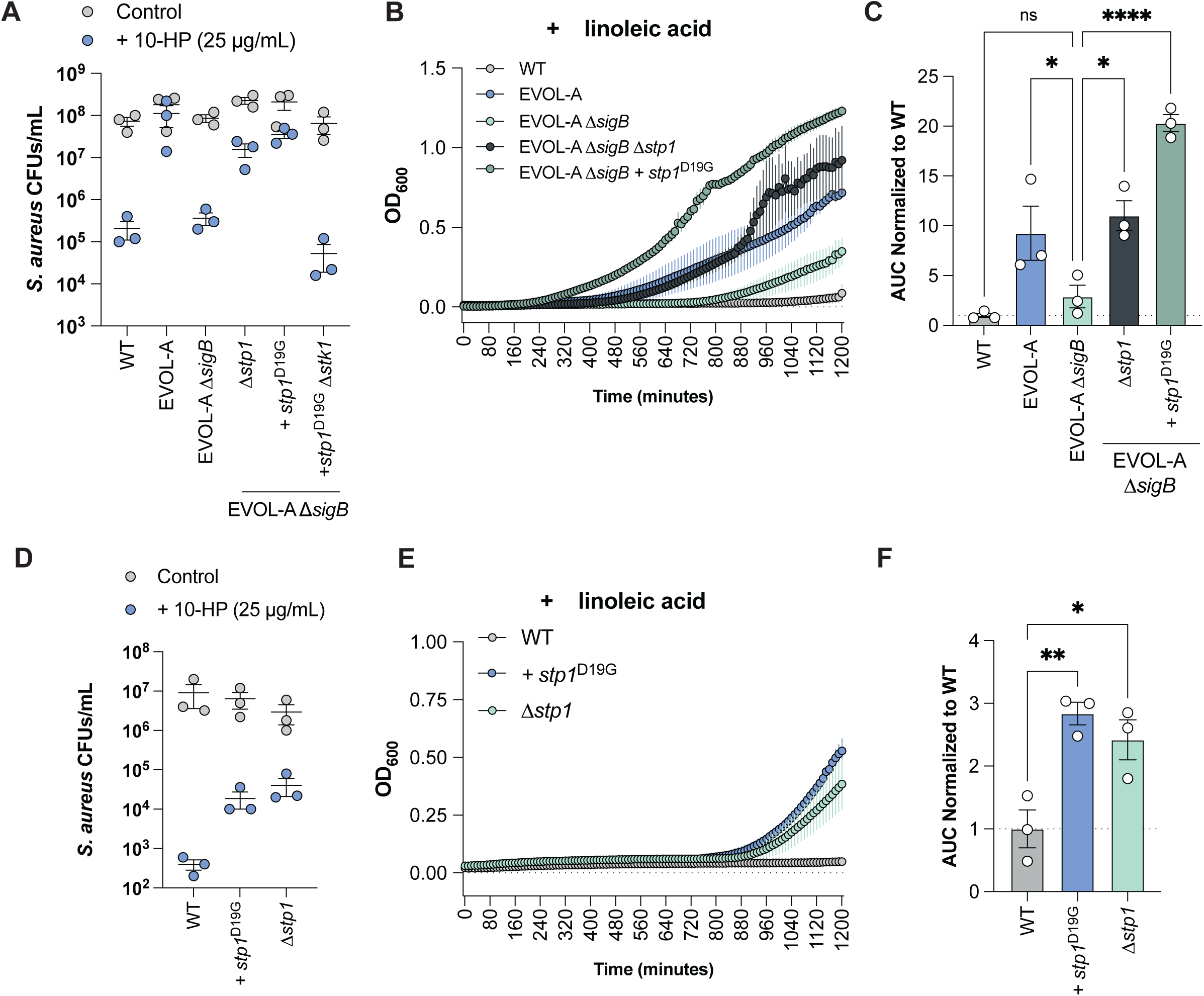
Stp1 loss of function mutation restores AFA tolerance in the absence of SigB. A. Recovered colony forming units (CFUs) after treatment with 10-HP or the control condition for 4 hours. Each point represents the mean of an independent experiment. B. 20-hour growth curve in TSB with 112 µg/mL linoleic acid showing mean OD_600_ from three separate experiments. C. AUC calculated from growth curves in A. Each point represents a separate experiment. D. Recovered colony forming units (CFUs) after treatment with 10-HP or the control condition for 4 hours. Each point represents the mean of an independent experiment. E. 20-hour growth curve in TSB with 112 µg/mL linoleic acid showing mean OD_600_ from three separate experiments. F. AUC calculated from growth curves in E. Each point represents a separate experiment. *:p<0.05, **:p<0.01, ***:p<0.001, ****p<0.0001, ns: p>0.05

We next sought to determine if loss of *stp1* was sufficient to establish AFA tolerance in the absence of the Rel truncation mutation. We introduced the loss of function allele (*stp1*^D19G^) and generated the *stp1* deletion (Δ*stp1*) in the WT (NRS193) background. Both the mutant allele and complete loss of *stp1* conferred increased tolerance to Ms-CFS (**Figure S4G**), 10-HP (**Figure 4D**), and resulted in significantly increased growth in the presence of linoleic acid (**Figure 4E, F**). However, the magnitude of the increased survival in 10-HP compared to WT was more modest for both +*stp1*^D19G^ and Δ*stp1* (45 to 100-fold, respectively) in the WT background compared to EVOL-A Δ*sigB*. This suggests that although *stp1* loss of function is sufficient to increase AFA tolerance, there may be other physiological changes conferred by the Rel mutation that alter AFA sensitivity.

### AFA tolerance coincides with phenotypic alteration of the cell envelope

Stk1/Stp1 mutations often confer cell wall changes and have been associated with tolerance to the cell wall targeting antibiotic vancomycin in vancomycin intermediate *S. aureus* (VISA) strains (50, 51). The enrichment for *walK* mutants, another gene implicated in the VISA phenotype and cell wall thickness, during EVOL-A Δ*sigB* adaptation to Ms-CFS further implicates cell wall changes in AFA tolerance (50). SigB overexpression has also previously been shown to increase the *S. aureus* cell wall thickness by ∼20% (52), and deletion of *sarA* in VISA strains has been shown to restore vancomycin susceptibility (53) Our previous work also described SigB-dependent vancomycin tolerance in EVOL-A (27). Together these data suggest that changes in the cell wall may occur downstream of Rel truncation to mediate AFA tolerance.

To test this, we first visualized the bacterial cell wall using transmission electron microscopy (TEM) in the WT, EVOL-A, EVOL-A Δ*sigB*, and EVOL-A Δ*sigB* +*stp1*^D19G^ strains (**Figure 5A**). It was immediately obvious that EVOL-A cells are smaller than WT and across non-dividing cells (without septa) cell diameter is significantly shorter (**Figure 5B**). Interestingly, the smaller cell size appeared to track with reduced AFA sensitivity, where cell diameter significantly increased in EVOL-A Δ*sigB* compared to EVOL-A and was restored to a shorter length with the introduction of *stp1*^D19G^. Despite overall smaller cell size, the cell wall thickness of EVOL-A is significantly greater than WT and EVOL-A Δ*sigB*, and the introduction of *stp1*^D19G^ in EVOL-A Δ*sigB* increases the cell wall thickness once again (**Figure 5C**). Most *S. aureus* strains have a cell wall that is 20-40 nm thick(54), and our WT NRS193 has an average cell wall thickness within this range at 29.9 +/-2.6 nm (**Figure 5A, B**). The thicker EVOL-A cell wall is on average 39.0 nm +/-2.8 nm thick and phenotypically distinct from WT, as it appears more compact and electron dense. EVOL-A Δ*sigB* cell wall thickness is similar to WT at 31.9 +/-2.6 nm and subsequent introduction of *stp1*^D19G^ restores cell wall thickness to EVOL-A levels (39.1 +/-2.8 nm). Together, these data indicate the AFA tolerant cells have reduced cell size with increased cell wall thickness.

**Figure 5.**
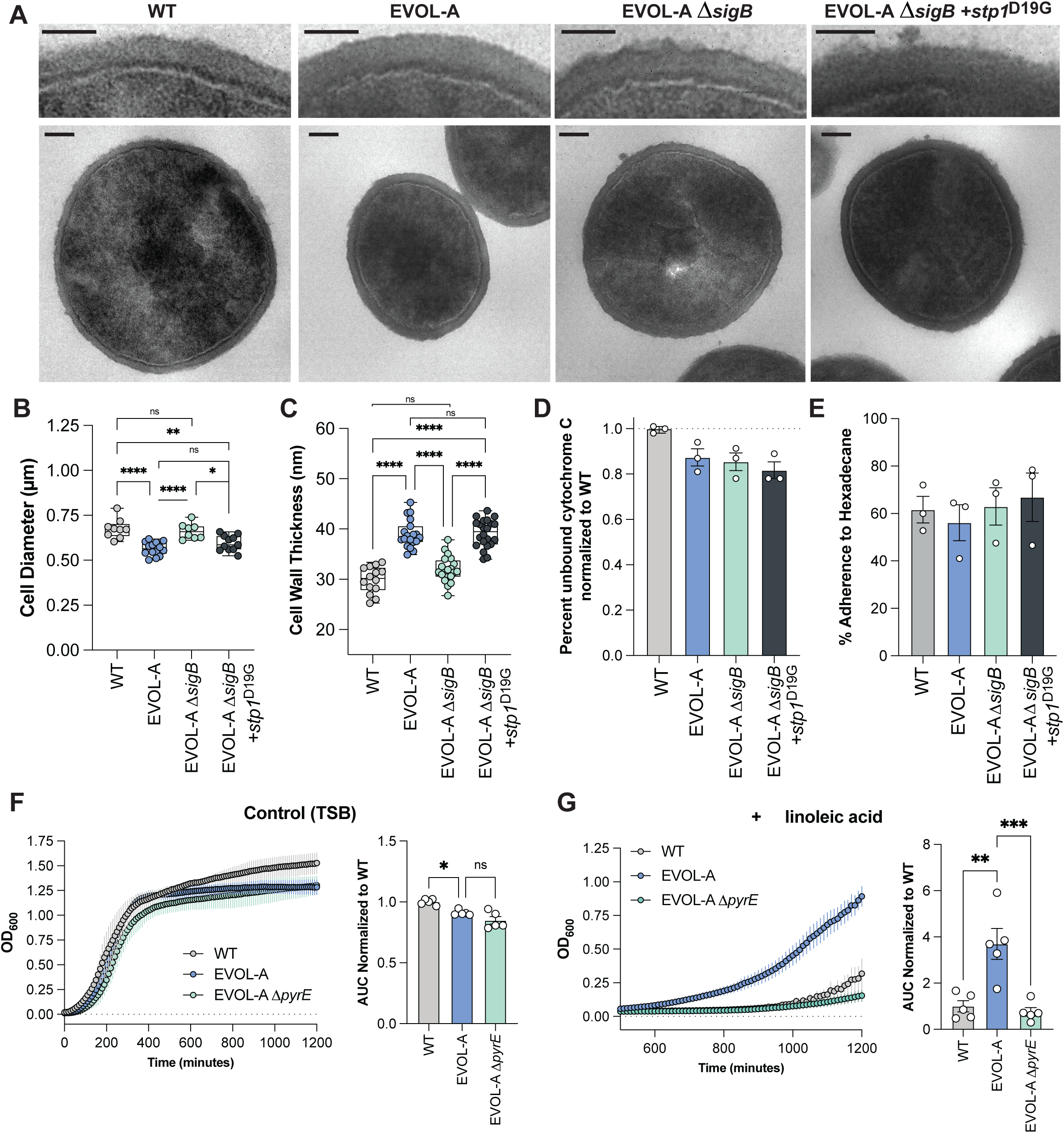
Stp1 loss of function restores the SigB-dependent thickened cell wall of the Rel mutant EVOL-A. A. Representative transmission electron micrographs of *S. aureus* strains grown in basal conditions. Top scale bars are 50 µm and bottom scale bars are 100 µm. B. Cell diameter quantified from transmission electron micrographs of non-dividing cells. C. Cell wall thickness quantified from transmission electron micrographs. D. Percent Cytochrome C remaining unbound after incubation with *S. aureus* strains normalized to WT. Each point represents an independent experiment. E. Percent affinity for the hydrophobic solvent n-hexadecane for *S. aureus* strains. Each point represents an independent experiment. F. 20-hour growth curve in TSB showing mean OD_600_ from five separate experiments with AUC calculated from growth curves. Each point on the bar graph represents a separate experiment. G. 20-hour growth curve in TSB with 112 µg/mL linoleic acid showing mean OD_600_ from five separate experiments with AUC calculated from growth curves. Each point on the bar graph represents a separate experiment. *:p<0.05, **:p<0.01, ***:p<0.001, ****p<0.0001, ns: p>0.05

The cell envelope, which includes the cell membrane and the cell wall, is well documented to influence AFA sensitivity in *S. aureus.* For example, wall teichoic acids and surface proteins impact cell surface change and hydrophobicity to protect cells from AFAs (10, 55). We sought to determine if such bulk envelope characteristics might be altered in AFA tolerant strains. We measured cell surface charge relative to WT through binding of the positively charged protein cytochrome C. While cell surface change was slightly reduced in EVOL-A compared to WT, EVOL-A Δ*sigB* was not different from EVOL-A (**Figure 5D**). This suggests that cell surface charge is not contributing to the AFA tolerance in these strains. We next assessed cell surface hydrophobicity by measuring the affinity of each strain for the hydrophobic solvent n-hexadecane. *S. aureus* cells with reduced surface hydrophobicity have been shown to evade AFA toxicity (10). However, there were no significant changes in affinity for n-hexadecane between WT, EVOL-A, EVOL-A Δ*sigB*, or EVOL-A Δ*sigB* + *stp1*^D19G^ (**Figure 5E**). This suggests that while the cell envelope is thicker in the AFA tolerant strains, this does not reflect changes in surface charge or hydrophobicity.

### SarA-dependent pyrimidine biosynthesis is necessary for AFA tolerance in a Rel truncation mutant

We next sought to understand the molecular mechanism by which the potential increased SigB regulation of SarA leads to AFA tolerance. In the EVOL-A strain, one of the clearest transcriptional signatures is a SarA-dependent increase in the expression of genes involved in pyrimidine biosynthesis (**Figure 3A, D**). Notably, Stk1/Stp1 have also been described to regulate pyrimidine biosynthesis in *S. aureus* and other Gram-positive bacteria (49). Biosynthesis of the pyrimidines uracil and cytosine is also essential for cell envelope homeostasis. Synthesis of uridine diphosphate *N*-acetylglucosamine (UDP-GlcNAc) is a key intermediate for peptidoglycan synthesis, and cytidine diphosphate diacylglycerol (CDP-DAG) is central to the synthesis of the membrane lipid phosphatidylglycerol (56, 57). A recent study of a Δ*pyrE* strain generated in the *S. aureus* USA300 LAC background reported the mutant to have a significantly thicker cell wall and irregular cell membrane supporting the role for pyrimidines in cell envelope homeostasis (58). To determine the contribution of pyrimidine metabolism to AFA sensitivity, we first leveraged the NTML *pyrF*::Tn strain and a low dose treatment of the AFA 10-HP. Compared to the WT control (JE2), a USA300 LAC derivative, interruption of the uridine monophosphate synthase (pyrF::tn) displayed increased sensitivity to 10-HP, with a ∼100-fold reduction in viable cells compared to 10-HP treated WT (**Figure S3B**). This result was somewhat unexpected, as pyrimidine limitation through an in-frame deletion of the orotate phosphoribosyltransferase (*pyrE*) was shown to increase *S. aureus* cell wall thickness in a related strain, and our data suggests a thicker cell wall is associated with AFA tolerance (58). However, in the previous study of *pyrE*, the Δ*pyrE* cells were grown in TSB and still displayed reduced growth compared to WT suggesting pyrimidine limitation. We grew the WT (JE2) and *pyrF*::tn in TSB for 20 hours and did not detect a difference in growth as measured by the area under the curve (AUC) (**Figure S3C**). This suggests that in contrast to the previous study of LAC Δ*pyrE*, in our assays the *pyrF*::tn strain is not experiencing pyrimidine limitation. Notably, when grown in the presence of 112 µg/mL linoleic acid, the *pyrF*::tn strain showed a significant reduction in growth compared to WT (**Figure S3C**). From these data we conclude that in pyrimidine replete conditions, the *de novo* pyrimidine biosynthesis pathway contributes to AFA tolerance in *S. aureus*.

We next sought to determine the contribution of pyrimidine biosynthesis to the increased AFA tolerance of the Rel mutant strain EVOL-A. To do so, we generated an in-frame Δ*pyrE* strain in the EVOL-A background, where *pyrE* is elevated in expression greater than 32-fold compared to the WT (NRS193) (**Table S2**). As reported previously, deletion of *pyrE* resulted in the accumulation of extracellular orotate crystals when grown on agar (data not shown)(58). Similar to the disruption of *pyrF* in the JE2 strain, deletion of *pyrE* in EVOL-A did not significantly impact growth in TSB compared to EVOL-A (**Figure 5F**). This indicates that TSB represents pyrimidine replete conditions in our assays. When grown in TSB with 112 µg/mL linoleic acid, loss of *pyrE* completely abolishes the enhanced growth of the EVOL-A strain restoring AFA sensitivity to WT levels (**Figure 5G**). Together this data suggests that even in pyrimidine replete conditions, the *de novo* pyrimidine pathway contributes to the AFA tolerance of the Rel truncated *S. aureus* strain EVOL-A. While the influence of pyrimidines on cell envelope homeostasis may be a contributing factor to AFA tolerance, further study is necessary to support this model, as disruption of pyrimidine biosynthesis has been shown to have large scale effects on *S. aureus* physiology (58). Collectively our work has identified a previously unappreciated role for the stringent response regulator Rel in AFA tolerance in *S. aureus* that arises through C-terminal truncation and a subsequent SarA-mediated regulatory cascade impacting the cell envelope.

## Discussion

The stringent response (SR) is a highly conserved stress response in bacteria(29). In *S. aureus,* the SR is mediated by the tightly regulated balance of alarmone production and degradation by the pyrophosphokinase Rel. Over production of alarmones completely stalls bacterial growth and the inability to activate the SR results in a rapid loss of viability during stress (33, 59, 60). Despite its highly conserved functions and conditional essentiality in *S. aureus*, mutations in Rel are being increasingly reported in clinical isolates and publicly available genomes (27, 34, 36). This supports a model that during adaptation to the host, Rel mutations tune SR activation to survive antibiotic exposure (35), antagonism by the microbiome (27), nutrient limitation (34), or to persist within host cells (61). Recent work also suggests a role for these mutations in controlling intraspecies transfer of plasmids (62).

In our previous work, we identified mutations throughout the Rel coding sequence that confer survival to *S. aureus* when exposed to the antimicrobial fatty acid (AFA) 10-HP produced by *M. sympodialis* (27). These Rel variants were derived from *in vitro* selection for survival following antagonism by *M. sympodialis* (lab variants) or were identified in *S. aureus* clinical isolates and publicly available genomes (natural variants) (**Figure 1A**). We were specifically interested in the C-terminal Rel truncation mutants that are not as well characterized as mutations in the N-terminal catalytic domain. Engineered truncation of the C-terminal ACT domain in Rel has been shown to cause (p)ppGpp overproduction by disrupting Rel’s normal interactions with stalled ribosomes, thereby dysregulating its catalytic activity (63). However, our previous measurements of (p)ppGpp in the Rel-Q672* strain, which truncates the final 57 amino acids, did not reveal alarmone levels different than WT after induction with mupirocin (27). Furthermore, 10-HP tolerance in this strain is independent of *codY* which responds to reduced GTP levels during (p)ppGpp production to de-repress transcription. Instead, the 10-HP tolerance is dependent on increased activity of the alternative sigma factor SigB (27).

The SR transcriptome has been studied extensively in *S. aureus* and includes significant changes in expression for over 100 genes (39, 64). In one such example, SR activation by induction of the Rel synthetase domain resulted in a significant increase in the SigB reporter gene *asp23* and a significant increase in expression of the staphylococcal accessory regulator *sarA* (64). SigB binds to the *sarA* P3 promoter to positively regulate transcription in many, though not all, *S. aureus* strains (44). Our data suggests that this induction of *sarA* by SigB occurs independent of measurable (p)ppGpp production, GTP depletion, and CodY de-repression. It remains an open question how the truncation of Rel in *S. aureus* leads to elevated SigB activity. In Gram-negative bacteria (p)ppGpp directly interferes with RNA-polymerase and sigma factor interactions selecting for associations with alternative sigma factors(65). However, this has not been shown to occur in *S. aureus.* Notably, high levels of active *sigB* are suggested to be toxic to *S. aureus*. The regulation of SigB by RsbU, RsbV, and RsbW provides multiple levels of control over SigB activity (66). Strains where this regulation is disrupted and SigB activity is unrestricted result in unstable small colony variants that spontaneously become SigB loss of function mutants (67). Perhaps mutations in Rel provide stable mechanisms for increased SigB activity with minimal toxicity to the cell. In *E. coli,* the Rel homolog SpoT interacts with Rsd, the RpoD (σ^70^) anti-σ activity protein, impacting SpoT hydrolase activity and stringent response activation, suggesting that C-terminal mutations in Rel could disrupt unknown protein-protein interactions that impact SigB activity (68).

In this work we have primarily focused on AFA tolerance mediated by the C-terminal Rel mutant Rel-Q672* (EVOL-A). While we have demonstrated that tolerance to 10-HP and linoleic acid are also mediated by *sigB* and *sarA* for an additional C-terminal mutant Rel-E657*(**Figure S2**), their contributions to AFA tolerance in other types of Rel mutants remain to be investigated. For the Rel hydrolase mutant Rel-F128Y, we have previously reported elevated SigB activity that is necessary for 10-HP tolerance, but the role of *sarA* in this strain is unknown. It is interesting that the hydrolase mutants Rel-F128Y and Rel-L127V appear to grow better than any of the C-terminal Rel mutants when exposed to the AFA linoleic acid (**Figure 2G, H**). This might suggest that these mutants either have even greater SigB and SarA activity, or that additional mechanisms of AFA tolerance are active in these strains.

Several mechanisms of AFA tolerance have been documented in *S. aureus*. These include expression of the fatty acid efflux pump FarE (26) detoxification of unsaturated fatty acids by the oleate hydratase OhyA (25), and detoxification by forming AFA-cholesterol esters via Lip2 (24). From our RNA-seq data comparing EVOL-A to WT at baseline conditions, *farE* and *isdA* are reduced in expression in EVOL-A and *ohyA* and *lip2* are unchanged in expression in EVOL-A (**Table S3**). While not definitive evidence that these AFA tolerance mechanisms are uninvolved, it suggests they are not playing a significant role in EVOL-A. The observation that cell surface charge and surface hydrophobicity are not changed in a SigB or Stp1-dependent manner also suggests that AFA tolerance is not occurring through alteration of wall teichoic acids in EVOL-A (**Figure 5 D, E**) (55).

Our current data instead supports a model where a thicker cell wall is associated with AFA tolerance. In support of this, VISA strains, which typically have thicker cell walls, were previously shown to have reduced sensitivity to *M. sympodialis* antagonism compared to other clinical *S. aureus* isolates (27, 50). Furthermore, in the absence of *sigB* and *sarA*, tolerance to *M. sympodialis* and 10-HP evolved independently through mutations in *walK* and *stp1*, two genes often mutated in VISA clinical isolates (**Table 2**) (50). The cell wall thickening observed in the Rel mutant EVOL-A compared to WT (∼34% increase) is modest compared to reported VISA strains (40-100% increase) (69, 70). Yet, the increase is similar to that reported for *sigB* overexpression (52). Cell wall remodeling and thickening can represent a non-specific adaptation to tolerate antimicrobials, where changes in the cell wall can broadly reduce antimicrobial penetration and access to targets. The specific targets of many AFAs, including 10-HP and linoleic acid, remain to be fully defined, but both AFAs can induce membrane damage (16, 19). Based on these data, we hypothesize that cell wall thickening and remodeling is one mechanism by which *S. aureus* can adapt to tolerate AFAs by preventing AFA access to the cell membrane. In addition to increased thickness, chemical and structural changes in the EVOL-A cell wall remain to be defined, but even by TEM the cell wall appears qualitatively different than WT (**Figure 5A**). It is also possible that the changes in the cell wall reflect changes in the cell membrane where cell wall biosynthetic machinery is anchored. Future studies will investigate changes in cell membrane characteristics including fluidity and polarization.

It remains an open question how the SarA-dependent increase in expression of pyrimidine biosynthesis genes may contribute to AFA tolerance. Pyrimidines are essential for cell envelope precursors. For example, UDP sugar derivatives, such as UDP-N-acetylglucosamine, are key cell wall building blocks that feed directly into peptidoglycan assembly (57). While we have observed that loss of *pyrE* in EVOL-A restores sensitivity to linoleic acid (**Figure 5G**), future work is needed to confirm whether this coincides with a reduction in cell wall thickness. Notably, recent work in *S. aureus* has reported that loss of *pyrE*, resulting in pyrimidine limitation, produces a thicker cell wall seemingly contrary to our hypothesis (58). Quantification of cell wall or cell envelope thickness from TEM overlooks underlying chemical and structural changes that may also mediate AFA tolerance. As such, changes in the cell wall that result from increase pyrimidine biosynthesis or pyrimidine limitation may be quite different but indistinguishable at the available resolution. Further studies are necessary to tease apart the role of the *de novo* pyrimidine biosynthesis pathway in cell envelope homeostasis.

In summary, this work reveals how Rel mutations in *S. aureus* affect AFA sensitivity. Even small Rel truncations markedly alter AFA sensitivity without canonical SR transcriptional signatures, instead acting through SigB-dependent regulation of SarA. Preliminary data suggest that increased cell envelope thickness may contribute to AFA tolerance, though the mechanism remains unclear. Because AFAs are abundant at host sites such as skin and blood these mutations may confer a niche-specific advantage. Elevated SarA activity downstream of Rel truncation raises questions about how *in vivo* Rel mutations influence virulence traits such as biofilm formation, toxin production, and immune evasion (71).

## Materials and Methods

### Microbial growth conditions

*S. aureus* strains were maintained on tryptic soy agar (TSA) or broth (TSB) at 37°C. Unless otherwise noted, *S. aureus* was cultured overnight to stationary phase (∼16-h) and sub-cultured 1:10 in TSB for 1 hour corresponding to early exponential phase. The wild type (WT) *S. aureus* in this study is NRS193 (ST1, USA400). NRS192 is closely related to the USA400 reference strain MW2. *E. coli* strains were grown in LB with antibiotic as needed: 100 mg/mL ampicillin or 25 mg/mL chloramphenicol.

### Bacterial growth curves

Growth curves were performed using a Synergy H1 monochromator-based multi-mode microplate reader (Agilent, Bio Tek). Non-treated, polystyrene 96-well plates with lids were utilized and growth was monitored by OD_600_ at 10-minute intervals over 20 hours. During the experiment, the plate was incubated at 37°C with continuous orbital shaking between measurements. Wells were inoculated at an initial OD_600_=0.02 in 150 µL of media. Each experiment included a minimum of three technical replicates. The untreated condition was TSB with vehicle control, which was ethanol for all three fatty acids. The fatty acids used were as follows: α-linoleic acid (Sigma-Aldrich, ≥98%) at 112 µg/mL, lauric acid (Cayman Chemical, ≥98%) at 75 µg/mL, and *cis-*palmitoleic acid (Sigma-Aldrich, ≥98.5%) at 37.5 µg/mL. Data was collected in Bio Tek Gen5 software and analyzed in Microsoft Office Excel and GraphPad Prism 10.

### RNA preparation, sequencing, and analysis

Samples for RNA-sequencing were prepared to determine baseline changes in transcription prior to any treatment. *S. aureus* NRS193, EVOL-A, or EVOL-A Δ*sarA* were cultured in TSB overnight for 16 hours at 37°C shaking at 250 rpm in three independent, biological replicates. After 16 hours, the cultures were back diluted 1:10 in fresh TSB and returned to 37°C shaking at 250 rpm for 1 hour. After the subculture, the 5 mL cultures all had an OD_600_ between 1.50 and 1.61. Cultures were centrifuged at 5000 rcf for 5 minutes at room temperature and cell pellets were resuspended in 1 mL of RNA Later (Ambion) for storage at 4°C for 24 hours.

To extract RNA, the preserved cells were centrifuged for 1 min. at 13,000 rcf and the pellets were resuspended in TE buffer containing 100 µg/mL lysostaphin (Sigma Aldrich). After 30 minutes at 37°C, 700 µL of RLT buffer with 2-mercaptoethanol from the Qiagen RNeasy Plus Mini Kit was added. The entire volume was transferred to a 2 mL tube with 0.1 mm disruption beads, and cells were pulsed 3 times for 30s with 10s rest using a Benchmark Bead Bug homogenizer. Samples were then centrifuged for 3 minutes at full speed and supernatants were transferred to the Qiagen gDNA eliminator column. The Qiagen RNeasy protocol were then followed as instructed. RNA quality was determined by fragment analysis and all samples had RQN greater than 7.9.

RNA processing, Illumina sequencing, and analysis were performed by SeqCenter (Pittsburgh, PA). Briefly, samples were DNAse treated (Invitrogen) and libraries were prepared by rRNA depletion using the Illumina Standard Total RNA Prep with Ribo-Zero plus Microbiome kit. Sequencing was performed on a NovaSeq X Plus generating 150 bp reads. RNA-sequencing data has been deposited in NCBI under GEO-TBD. The reference genome for NRS193 was provided for gene annotation and analysis (NCBI SAMN41193238) (27). Adapter trimming and quality control were performed using Illumina’s bcl-convert (version 4.2.4) with default parameters. Read mapping was performed with HISAT2 (version 2.2.1) using default parameters+’--very-sensitive’ (72). Read quantification was performed using Subread’s featureCounts (version 2.0.6) using default parameters + ‘-Q 20’ (73). Reads were normalized in R (version 4.0.2) using edgeR’s Trimmed Mean of M values (TMM) algorithm with default parameters and values were converted to counts per million (CPM) (74, 75). Analysis of differential expression was performed using edgeR’s glmQLFTest and a subset of genes with |log_2_FC| > 1 and p <.05 were determined to have significant differential expression between compared groups. **Table S3** and **Table S5** show the CPM and edgeR glmQLFTest results for all genes.

### S. aureus allelic exchange

Allelic exchange in *S. aureus* was performed using methods previously described (27, 76). All plasmids and primers are provided in **Table S6**. For generation of in frame gene deletions, the pIMAY system was used. Briefly, knock out constructs were generated by amplifying 500-1000 bp 5’ and 3’ of the gene of interest. Overlap extension PCR was performed to fuse the two fragments, then the product was introduced into SmaI digested pIMAY, and transformed in *E. coli* DC10B. Colonies were isolated on TSA with 25 µg/mL chloramphenicol. PCR confirmed plasmids were electroporated into *S. aureus* and colonies were selected for on TSA with 10 µg/mL chloramphenicol at 28°C. Large colonies were then resuspended and plated on TSA with 10 µg/mL chloramphenicol to select for chromosomal integration. PCR confirmed colonies with pIMAY recombination on the chromosome were then grown overnight at 28°C in the absence of chloramphenicol and then plated on TSA with 1 µg/mL anhydrotetracycline hydrochloride (aTC) at 28°C for 24-48 hours. Single colonies were patched on TSA with 1 µg/mL aTC and TSA with 10 µg/mL chloramphenicol. Colonies that grew on aTC but not chloramphenicol were confirmed by PCR to have the desired gene deletion.

For the generation of PsarA-YFP, the entire promoter region for sarA (P1, P2, P3) was amplified using primers PsarA_F and PsarA_R (**Table S6**). The product was column purified and digested with SbfI and SmaI. The previously published Pasp23-YFP plasmid (pIMAY derived pKOW1) was digested with SbfI and SmaI to remove the asp23 promoter upstream of GPVenus (YFP). The digested sarA promoter was ligated into the digested pKOW1 using NEB’s Quick Ligation Kit before being transformed into *E. coli* DC10B. pKOW1 contains flanking regions to target chromosomal insertion of the transcriptional reporter to a previously determined neutral site (43). Transformation into *S. aureus* and recombination for pIMAY derived plasmids was performed as described above (27, 76).

For the *stp1* allele swap, primers were designed to amplify a single fragment from the mutant genome including 1000 bp on either side of the mutation (D19G) (Stp1_swap_F and Stp1_swap_R) (**Table S6**). The resulting DNA fragment was introduced into SmaI-digested pIMAY and transformed into *E. coli* DC10B. The transformation of *S. aureus* was then performed as described above for pIMAY.

### M. sympodialis CFS preparation and 10-HP tolerance assays

*M. sympodialis* strain KS269 was maintained in mDixon media and cultured at 30°C. mDixon was prepared as follows: 36 g/L malt extract, 20 g/L ox-bile, 10 mL/L tween40, 6 g/L peptone (from casein and other animal proteins), 2 mL/L glycerol, 2 mL/L oleic acid, 15 g/L agar, and adjusted to pH 6 with HCl. The preparation of *M. sympodialis* cell-free supernatant was prepared as previously described (27). Briefly, *M. sympodialis* was inoculated into mDixon broth and cultured at 30°C for 96 hours shaking at 200 rpm. Cultures were centrifuged at 5000 rcf for 2 minutes and the supernatant was decanted. The pH of the supernatant was adjusted to pH 5.5 with NaOH and/or HCl. The supernatant was then filter-sterilized with a 0.22 µm MCE filter to generate the *M. sympodialis* CFS (Ms-CFS).

For experiments where *S. aureus* was treated with Ms-CFS, a concentration of 50% was generated by mixing Ms-CFS one-to-one with pH matched mDixon broth. *S. aureus* cells were cultured overnight and sub-cultured 1:10 for 1 hour as described above in TSB. Cells were then pelleted and resuspended in mDixon and inoculated into 50% Ms-CFS or pH matched mDixon broth at an initial OD_600_ of 0.02. Treatment times are described in the figure legends. After treatment *S. aureus* viability was enumerated by plating for CFUs on TSA.

For 10-HP treatment assays, 10-HP (Ambeed) was resuspended in DMSO and stored at-80°C. 10-HP was added to mDixon at concentrations provided in the text and figure legends: 25 µg/mL (normal dose) or 10 µg/mL (low dose). Treatment time is also described in figure legends and ranges from 2-4 hours. After treatment, *S. aureus* viability was enumerated by counting CFUs plated on TSA after serial dilution.

### Experimental evolution in M. sympodialis CFS

The experimental evolution of *S. aureus* USA200 MN8 in Ms-CFS was performed as previously described (27). Briefly, MN8 was treated in three replicate populations of 50% Ms-CFS and one population of pH-matched mDixon broth control for 8 hours at 37°C with a starting inoculum of OD_600_=0.02. After 8 hours, 10 µL was removed to enumerate viability by plating for CFUs. The remaining 190 µL was inoculated into 5 mL of TSB to recover surviving cells overnight. This process was repeated for 12 passages. From the viable counts after passage 12, individual colonies were selected to assess Ms-CFS tolerance.

The experimental evolution assays of *S. aureus* EVOL-A Δ*sigB* and EVOL-A Δ*sarA* were carried out similar to the above experiment with a few alterations. The EVOL-A Δ*sigB* experiment used 50% Ms-CFS exposure for 4 hours and was repeated for only 6 passages. The EVOL-A Δ*sarA* experiment used 50% Ms-CFS exposure for 2 hours and was repeated for only 5 passages. These changes were made due to slight increases in Ms-CFS sensitivity in the mutant strains compared to NRS193 or MN8.

### DNA extraction, sequencing, and variant calling

*S. aureus* genomic DNA was extracted as previously described using the Qiagen DNeasy Blood and Tissue kit with modifications (27). *S. aureus* colonies from a fresh TSA plate were resuspended in TE buffer with 20 µg/mL lysostaphin (Sigma-Aldrich) and incubated at 37°C for 30 minutes or until visibly cleared. The lysis buffer for Gram-positive bacteria described in the DNeasy Blood and Tissue kit was prepared 2X without lysozyme and then diluted to 1X in the TE +lysostaphin. The Qiagen kit protocol was then followed as instructed.

Genome sequencing of the evolved strains was carried out by SeqCenter (Pittsburgh, PA) using Illumina whole-genome sequencing. Genomic DNA was prepared as described above, and libraries were constructed with the Illumina DNA Prep kit using bead-based tagmentation. The resulting Illumina sequencing reads (FASTQ files) were aligned to the NRS193 reference genome, and variants were identified with breseq using its default parameters (77).

### Transmission electron microscopy

Bacterial cultures were grown overnight, back-diluted to an OD600 of 0.05 and grown to OD600 0.5. The cultures were fixed with a 2.5% w/w glutaraldehyde in 0.1 M cacodylate buffer [pH 7.4] overnight at 4°C. The fixed samples were treated with 2% w/w aqueous osmium tetroxide for 2 hours at room temperature. Samples were dehydrated through a graded series of ethanol in water (75% v/v, 95% v/v, 100% v/v), cleared in propylene oxide, embedded in Epon resin and cured in an oven for 72 hours at 60°C. Ultrathin sections, approximately 85 nm thick, were sectioned using a Reichert Ultracut E ultramicrotome with a diamond knife and collected onto 200 mesh nickel grids. These were stained for 25 minutes with 3% w/w aqueous uranyl acetate, washed with deionized water, stained with Reynold’s lead citrate for 5 minutes, washed with deionized water and blotted dry. Sections were viewed on a Philips CM100 BioTWIN Transmission Electron Microscope at 100 kV fitted with a Gatan MultiScan 794 CCD camera.

### Cell diameter and cell wall thickness measurements

Cell diameter was quantified from TEMs in FIJI/ImageJ (78). Cell diameter was only quantified from cells that had no septum formation. A minimum of two measurements were taken for each cell and averaged. The size in pixels quantized through FIJI was converted to µm based on the scale bar to pixel conversion of each individual image. Similarly for the cell wall thickness measurements, FIJI was used to quantify the cell wall from the electron dense membrane to the outer most area of the wall. Four to five measurements were taken from a single cell around the entire circumference of the cell and averaged. Pixel quantification was converted to nm based on scale bar to pixel conversion.

### Cytochrome C Binding Assay

The cytochrome C binding assay was carried out as previously described (79). As described above, cells were cultured overnight in TSB at 37°C before a 1:10 subculture was performed for 1 hour in TSB. The OD_600_ was adjusted to 1.1 and 2 mL of cells were washed twice in 20 mM sodium acetate buffer. After two washes, the cells were resuspended in 500 µL 20 mM sodium acetate buffer with 25 µg/mL cytochrome C. Cells were incubated at 37°C with agitation for 15 minutes. Cells were then pelleted by centrifugation at 16,000 rcf for 2 minutes at room temperature. From the supernatant, 100 µL was transferred to a flat bottom 96-well plate. A control of buffer with cytochrome C but without cells was included. Absorbance was read at 410 nm. To calculate unbound cytochrome C, values were normalized to the no cell control.

### Microbial Adhesion to Hydrocarbon Assay

Cells were cultured overnight (∼16 hours) in TSB at 37°C before a 1:10 subculture was performed in TSB for 1 hour as described above. Cells were prepared for the Microbial Adhesion to Hydrocarbon Assay as previously described (80). Briefly, cell were centrifuged at 5,000 rcf for 2 minutes and washed three times with ice cold PBS. The OD_600_ was adjusted to 0.5 in glass tubes and 0.2 volumes of n-hexadecane was added. Tubes were vortexed for 1 minute and then left stationary at room temperature for 1 hour. The OD_600_ of the aqueous phase was measured and the percent adherence to hexadecane was calculated using the following equation: % adherence=(1-Abs/Abs_0_) x100. Where Abs_0_ is the initial OD_600_ of 0.5 and the Abs is the final OD_600_ of the aqueous phase.

### Statistics

Statistical analyses were performed in GraphPad Prism 10 (v. 10.4.1). All error bars indicate standard error of the mean (SEM). For comparisons between two groups, student’s t-test was used. For comparisons between two or more groups, a One-way ANOVA was performed with a Dunnett’s multiple comparisons test, when comparing all groups to a control, or a Tukey’s multiple comparisons test, when all groups were compared to each other. P-values calculated as >0.5 were determined to be not significant. Area under the curve for growth curve data was calculated in GraphPad Prism 10 with a baseline of Y=0, a min. peak height of 10% the distance from min. to max. Y, and all peaks must exceed the baseline. For all curves, only one peak was detected and quantified.

## Acknowledgments

We thank the University of Oregon Summer Undergraduate Research Experience and Bryan Rebar for their support of SBK in this work. Support for this work in the form of funding was received by CHK (L’Oréal USA for Women in Science Fellowship and Helen Hay Whitney Foundation Fellowship), SBK (UO VPRI Mini Grant), TJS (National Institutes of Health grants 1P01GM125576, F32DK124033, and K99DK137017), KC (National Institutes of Health T32GM149387), MFB (National Institutes of Health grants R35GM133652, R35GM158176, and R21AI173839), RMC (Sir Henry Dale Fellowship jointly funded by the Wellcome Trust and the Royal Society, 104110/Z/14/A, and a Lister Institute Research Prize Fellowship 2018).

## Supplemental Figure Legends

**Figure S1. Isolation of USA200 Rel mutant from experimental adaptation to Ms-CFS.**

A. Experimental schematic for serial passaging of *S. aureus* in Ms-CFS to select for Ms-CFS/10-HP tolerant strains.

B. Recovered USA200 colony forming units (CFUs) after exposure (passages) to Ms-CFS in three separate populations. Control population was exposed to pH-matched media control.

C. Recovery of a single isolate from USA200 Ms-CFS R2 passage 12 (P12) with a Rel mutation R233P after 2-hour treatment with 50% Ms-CFS. Data pooled from three experiments.

D. Recovery of a single isolate from USA200 Ms-CFS R2 P12 with a Rel mutation R233P after 2-hour treatment with 25 µg/mL 10-HP. Each point represents a separate biological replicate.

**Figure S2. Contribution of SigB and SarA to AFA tolerance in Rel-E657*.**

A. Recovered colony forming units (CFUs) after treatment with 10-HP or the control condition for 2 hours. Each point represents a separate biological replicate.

B. 20-hour growth curve in TSB with 112 µg/mL linoleic acid showing mean OD_600_ from four or six separate experiments depending on the strain.

C. AUC calculated from growth curves in B. Each point represents a separate experiment. *:p<0.05, **:p<0.01, ***:p<0.001, ****p<0.0001, ns: p>0.05

**Figures S3. AFA sensitivity of the NTML pyrF::Tn mutant.**

A. Recovered colony forming units (CFUs) after treatment with 25 µg/mL 10-HP or the control condition for 2 hours. Data pooled from two experiments.

B. Recovered colony forming units (CFUs) after treatment with 10 µg/mL 10-HP or the control condition for 2 hours. Data pooled from two experiments.

C. AUC calculated from 20-hour growth curves in TSB with or without 112 µg/mL linoleic acid. Data combined from two separate experiments. Students t-test performed. *:p<0.05, **:p<0.01, ***:p<0.001, ****p<0.0001, ns: p>0.05

**Figure S4. Tolerance to *M. sympodialis* antagonism evolves rapidly in the absence of *sigB* and *sarA*.**

A. Schematic of experimental evolution approach to select for 10-HP tolerance in EVOL-A Δ*sigB* through repeated exposure to Ms-CFS.

B. EVOL-A Δ*sigB* survival in Ms-CFS throughout the evolution experiment in three separate populations.

C. Ms-CFS tolerance of isolated strains from EVOL-A Δ*sigB* after six passages in Ms-CFS.

D. Schematic of experimental evolution approach to select for 10-HP tolerance in EVOL-A Δ*sarA* through repeated exposure to Ms-CFS.

E. EVOL-A Δ*sarA* survival in Ms-CFS throughout the evolution experiment in three separate populations.

F. Ms-CFS tolerance of isolated strains from EVOL-A Δ*sigB* after five passages in Ms-CFS.

G. Recovered CFUs after 2-hour treatment with 50% Ms-CFS or pH-matched media control. Pooled data from two experiments.

## Supplemental Tables

Table S1. Strains Used In This Study

Table S2. WT v. EVOL-A RNAseq Significant Differentially Expressed Genes

Table S3. WT v. EVOL-A RNAseq All Quantified Genes

Table S4. EVOL-A v. EVOL-A Δ*sarA* RNAseq Significant Differentially Expressed Genes

Table S5. EVOL-A v. EVOL-A Δ*sarA* RNAseq All Quantified Genes

Table S6. Primers And Plasmids Used In This Study

